# Over-expression of the Regulator of the Signaling of G Protein-Coupled Receptors RGS9-2 reduces the signaling elicited by the human histamine H_3_ receptor in HEK-293T cells

**DOI:** 10.1101/2021.03.22.435676

**Authors:** Gustavo Nieto-Alamilla, Juan Escamilla-Sánchez, José-Antonio Arias-Montaño

## Abstract

In HEK-293T cells transiently transfected with the human histamine H_3_ receptor (hH_3_R), we studied the effect of over-expressing the human RGS9-2 protein on H_3_R-mediated stimulation of [^35^S]-GTPγS binding and inhibition of forskolin-induced cAMP formation. Maximal specific binding (Bmax) of [^3^H]-N-methyl-histamine to cell membranes was 468 ± 12 and 442 ± 38 fmol/mg protein for HEK-293T-hH_3_R and HEK-293T-hH_3_R/hRGS9-2 cells, respectively, with dissociation constants (Kd) 2.57 nM and 3.38 nM. The H_3_R agonist immepip stimulated [^35^S]-GTPγS binding with similar potency and efficacy (E_max_ 146.3 ± 4.4 % and 150.0 ± 5.3 % of basal, pEC_50_ 8.57 ± 0.26 and 9.00 ± 0.33, respectively), but was significantly less efficacious to inhibit forskolin-induced cAMP accumulation in HEK-293T-hH_3_R/hRGS9 cells (−19.2 ± 5.3 versus −37.7 ± 5.1 % in HEK-293T-hH_3_R cells) with no significant difference in potency (pEC_50_ 9.60 ± 0.14 and 9.07 ± 0.29, respectively). These results indicate that in HEK-293T cells hRGS9-2 regulates hH_3_R_445_ signaling downstream G protein activation.

## 1. Introduction

In mammals, four G protein-coupled receptors (GPCRs) for histamine have been identified. Histamine H_3_ receptors (H_3_Rs) are abundantly expressed in the Central Nervous System (CNS), both pre- and post-synaptically. H_3_Rs couple to Gα_i/o_ proteins, and thereby inhibit adenylyl cyclases via Gα _i/o_ subunits, whereas the Gβγ complexes hinder the opening of N- and P/Q-type voltage-operated Ca^2+^ channels, resulting in inhibition of neurotransmitter release. H_3_R activation results in further actions, such as the opening of G protein-gated inwardly rectifying K^+^ channels (GIRKs), stimulation of phospholipases C and A_2_, and activation of the phosphatidylinositol 3-kinase (PI3K) and mitogen-activated protein kinase (MAPK) pathways [1–3].

The regulators of GPCR signaling (RGS) are proteins that enhance the GTPase activity of Gα subunits and modulate thus the intracellular effects elicited by GPCR stimulation and subsequent G protein activation. There exist 20 classical RGS proteins, classified into four subfamilies (RZ, R4, R7 and R12) based on sequence and domain homology [4,5]. Some RGS proteins co-express with H_3_Rs in cerebral cortex, hypothalamus, striatum and nucleus *accumbens* [6]. In the striatum, the isoform 2 of RGS9 (RGS9-2) is abundantly expressed, whereas the isoform 1 is expressed only in cone and rods of the retina cells. RGS9-2 enhances the GTPase activity of proteins of the Gα_i/o_ subfamily [7], and regulates the signaling exerted by dopamine and opioid receptors in striatal neurons [8,9].

To the best of our knowledge the possible modulation by RGS9-2 of H_3_R signaling has not been investigated. In this work we studied the effect of over-expressing the human RGS9-2 (hRGS9-2) on the activation of G proteins and signaling induced by the stimulation of the co-transfected human H_3_R of 445 amino acids (hH_3_R), the originally cloned receptor and the most abundantly expressed in the human brain along an isoform of 365 amino acids [10]. These experiments were performed in human embryonic kidney HEK-293T cells, widely used as a model to address RGS9-2 function [4].

## 2. Materials and methods

### 2.1 Cell culture and transfection

Parental HEK-293T cells (American Type Culture Collection) were cultured in Dulbecco’s modified Eagle’s medium (DMEM) with high glucose (1:1 v/v, Gibco, Thermo Fisher Scientific, Waltham, MA, USA) supplemented with 10 % fetal bovine serum, 2 mM glutamine, 0.01 mg/ml streptomycin and 100 U/ml penicillin. Cells were grown as monolayers at 37 °C in a CO_2_ incubator (5 % CO_2_).

Cell transfection was performed with lineal polyethylenimine (MW 25 kDa; Polysciences, Warrington, PA, USA). Parenteral HEK-293T cells (8×10^6^ cells) were seeded in Petri dishes (10 cm diameter) and grown by 24 h. DNA:PEI complexes were prepared in Eppendorf tubes containing 18 μg of total DNA (9 μg pCI-Neo-hH_3_R plus 9 μg pc-DNA3.1 or 9 μg pC-DNA3.1-RGS9-2) and 54 μl of sterile PEI (5 mg/ml). The volume was raised to ~1.8 ml with DMEM and then incubated at 37 °C for 30 min in a CO_2_ incubator. Cells were rinsed with sterile phosphate-buffered saline (PBS), 1.7 ml DMEM were added and the DNA:PEI complexes were then incorporated. After 30 min at 37 °C in a CO_2_ incubator, 3.5 ml DMEM (20% fetal bovine serum, 4 mM glutamine and 2% antibiotics) were added and cells were placed in the CO_2_ incubator (5 % CO_2_). After 15 h the medium was replaced by fresh complemented medium and the incubation continued for a total 48 h.

### 2.2 End-point RT-PCR

Total mRNA was extracted using a two-step Trizol reagent (Life Technologies). Samples (1 μg) were treated with DNAse I (Life Technologies), reverse-transcribed with SuperScript II Reverse Transcriptase First-Strand cDNA Synthesis System (Invitrogen, Carlsbad, CA, USA) and amplified for 28 cycles of 96 °C for 30 s, 60° C for 25 s, 72 °C for 30 s, and final extension at 72 °C for 7 min, using 1 U of Platinum Taq High Fidelity DNA polymerase (Invitrogen). The sequences of the primers directed to the third intracellular loop of the hH_3_R were: forward 5’-TCTTTACGCCCTTCCTCAGC-3’; reverse 5’-TCATCAGCAGCGTGTATGGG-3’, with expected fragments of 516 bp. The sequences of the primers used to detect the hRGS9-2 were: forward 5’-CCGCACAAACCCACATTTAC-3’; reverse 5’-GAAACCTGCTGAAGGACAGA-3’, with expected fragments of 588 bp. The products were analyzed in agarose gels (0.8 %) stained with ethidium bromide, and the relative expression levels of mRNAs were quantified using as reference the mRNA of the human ribosomal protein P0 (hRPLP0) amplified with the following primers: forward 5′-GCAGGTGTTTGACAATGGCAGC-3′; reverse 5′-GCCTTGACCTTTTCAGCAAGTGG-3′).

### 2.3 [^3^H]-NMHA binding to cell membranes

Parental and transfected HEK-293T cells, grown in plastic Petri dishes (10 cm diameter), were scrapped and homogenized in ice-cold lysis buffer (10 mM Tris-HCl, 1 mM EGTA, pH 7.4). Cell lysates were centrifuged (32,000xg, 20 min at 4°C) and the pellets re-suspended in incubation buffer (50 mM Tris-HCl, 5 mM MgCl_2_, pH 7.4). Saturating [^3^H]-NMHA binding (0.1-10 nM) was assayed in a 100 μl-final-volume with ~20 μg membrane protein (BCA assay; Pierce, Rockford, IL, USA), as described in detail elsewhere [11].

### 2.4 Western blotting

Assays were performed essentially as described previously [11]. Antibodies, dilutions and sources were as follows: rabbit anti hH_3_R (1:1000; Abcam EPR5631 or ab124732); rabbit anti human RGS9-2 (1:1000; Abcam EPR2873 or ab108975); goat anti-rabbit IgG (1:2000; Thermo Fisher Scientific cat. num.: 65-6120); anti-actin (1:1000; from hybridoma of a mouse cell line, kindly donated by Dr. José-Manuel Hernández-Hernández, Cinvestav); goat anti-mouse IgG (1:5000; Thermo Fisher Scientific cat. num.: 31430).

### 2.6 [^35^S]-GTPγS binding and cAMP accumulation assays

[^35^S]-GTPγS binding to cell membranes and cAMP accumulation in intact cells were assayed as described in detail elsewhere [11].

### 2.7 Drugs

The following drugs were purchased from Sigma Aldrich (Saint Louis, MO, USA): cAMP (adenosine 3,5-cyclic monophosphate), GTPγS (guanosine 5-[γ-thio]-triphosphate tetralithium salt), histamine dihydrochloride, IBMX (3-isobutyl-1-methylxantine) and immepip dihydrobromide. N-α-[methyl-^3^H]-histamine (85.4 Ci/mmol), [^35^S]-GTPγS (1250 Ci/mmol) and [5,8-^3^H]-adenosine 3,5-cyclic phosphate (34.0 Ci/mmol) were from Perkin Elmer (Boston, MA, USA).

## 3. Results

### 3.1 Expression of hH_3_R and hRGS9-2 by HEK-293T cells

Parental HEK-293T cells expressed low levels of endogenous hH_3_R mRNA (3.3 ± 0.4 % of the mRNA of hRPLP0, mean ± standard error, SEM; Figure 1A), and transfection with pCI-Neo-hH_3_R increased by 12.6 fold the levels of hH_3_R mRNA (4 determinations). The co-transfection of hRGS9-2 did not affect the expression of the hH_3_R mRNA (Figure 1A’).

**Figure 1.**
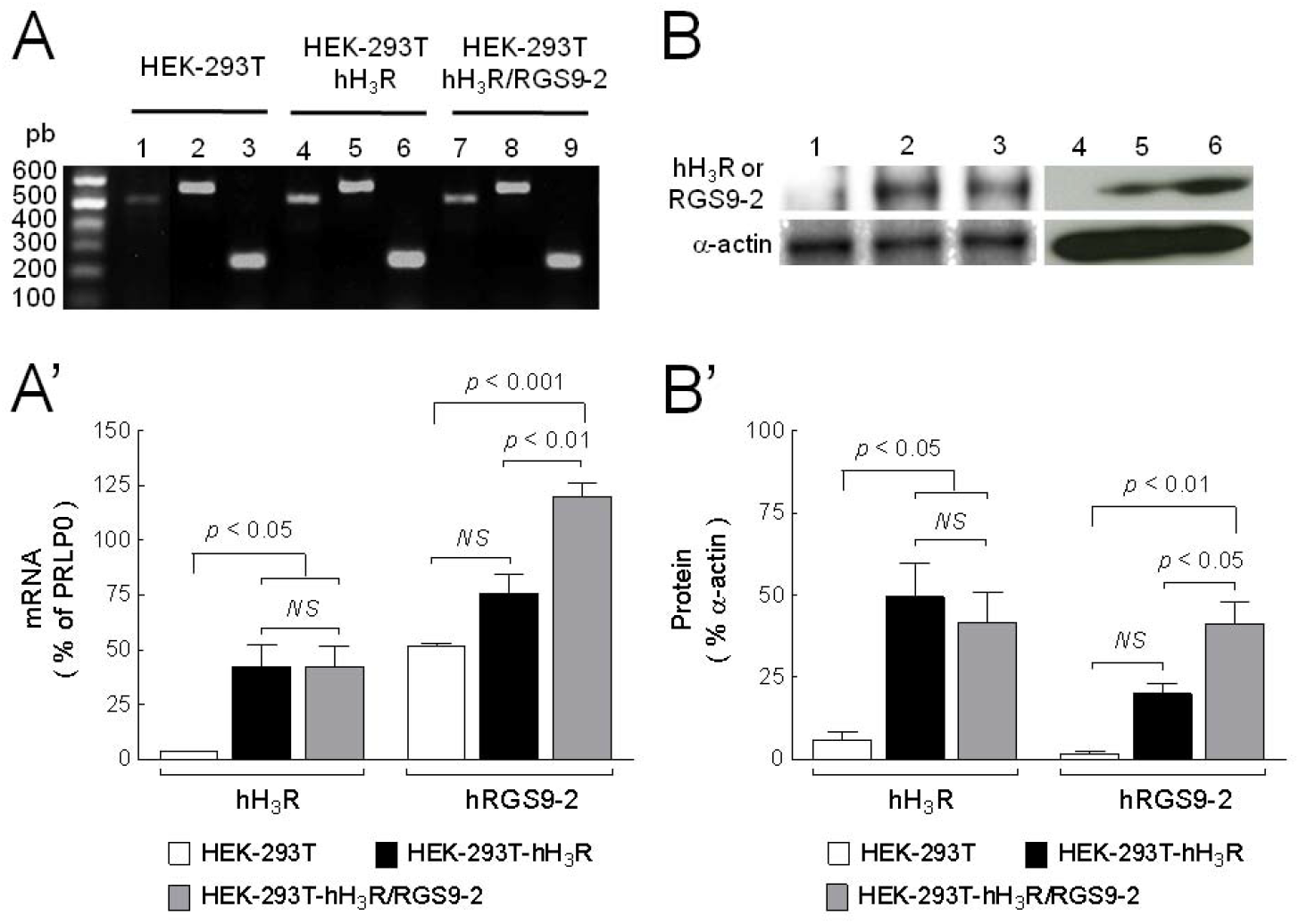
Expression of the hH_3_R and hRGS9-2 in transiently transfected HEK-293T cells. A. End-point RT-PCR. The amplicons were of 512 bp for the hH_3_R (lanes 1, 4 and 7), 588 bp for the hRGS9-2 (lanes 2, 5 and 8) and 231 bp for the endogenous hRPLP0 protein (lanes 3, 6 and 9) in parental, HEK293-hH_3_R and HEK293T-hH_3_R/hRGS9-2 cells. A representative determination is depicted. A’. Analysis of 4 experiments. Values are means ± SEM of the mRNA levels normalized to hRPLP0 transcripts. The statistical analysis was performed with ANOVA and Tukey’s test. B. Western blot analysis of hH_3_R and hRGS9-2 protein levels in parental, HEK293-hH_3_R or HEK293T-hH_3_R/hRGS9-2 cells. B. Blots from a representative determination show the hH_3_R (lanes 1-3) and hRGS9-2 proteins (lanes 4-6) in parental cells (lanes 1 and 4), HEK293-hH_3_R cells (lanes 2 and 5), and HEK293T-hH_3_R/hRGS9-2 cells (lanes 3 and 6). B’. Analysis of 6 experiments. Values are means ± SEM of the protein levels normalized to α-actin contents (ANOVA and Tukey’s test).

The mRNA of endogenous hRGS9-2 was 51.6 ± 1.4 % of hRPLP0 in parental HEK-293T cells and increased by 1.46 fold in cells transfected only with hH_3_R, although the values were not statistically different (Figure 1A). The co-transfection of hRGS9-2 and hH_3_R resulted in a significant increase in hRGS9-2 mRNA in comparison with both parental and HEK-293T- hH_3_R cells (2.31 and 1.57 fold, respectively; Figure 1A’).

Western blot analysis (Figure 1B’) showed that the levels of the endogenous hH_3_R protein in parental HEK-293T cells (Figure 1B’; 5.7 ± 2.6 % of the corresponding content of α-actin) were significantly increased in HEK-293T-hH_3_R and HEK-293T-hH_3_R/hRGS9-2 cells (8.89 and 7.50 fold, respectively), with no significant difference between the transfected cells.

Figure 1B’ shows that parental HEK-293T cells expressed low levels of the endogenous hRGS9-2 protein (1.7 ± 0.7 % of the α-actin content) and that transfection of hH_3_R or hH_3_R/hRGS9-2 increased the expression to 20.0 ± 2.9 % and 41.3 ± 6.8 %, respectively. Although the transfection of hH_3_R augmented the endogenous expression of hRGS9-2, the levels were significantly higher in cells transfected with the plasmid that encode the hRGS9-2 protein (Figure 1B’).

The expression of the hH_3_R in cell membranes was analyzed by [^3^H]-NMHA binding. Low [^3^H]-NMHA binding was detected in parental cells (7.1 ± 3.4 fmol/mg protein, 3 determinations). Saturation binding (Figure 2) shows that the transfection increased the density (B_max_) of the hH_3_R to 468 ± 12 fmol/mg protein, and that the over-expression of RGS9-2 did not affect the receptor levels (442 ± 38 fmol/mg protein, 4-5 determinations; *p* = 0.5108, Student’s *t* test) or the equilibrium dissociation constant (*K*_d_), 2.56 and 3.33 nM, respectively (p*K*_d_ 8.59 ± 0.04 and 8.48 ± 0.08, *p* = 0.2474).

**Figure 2.**
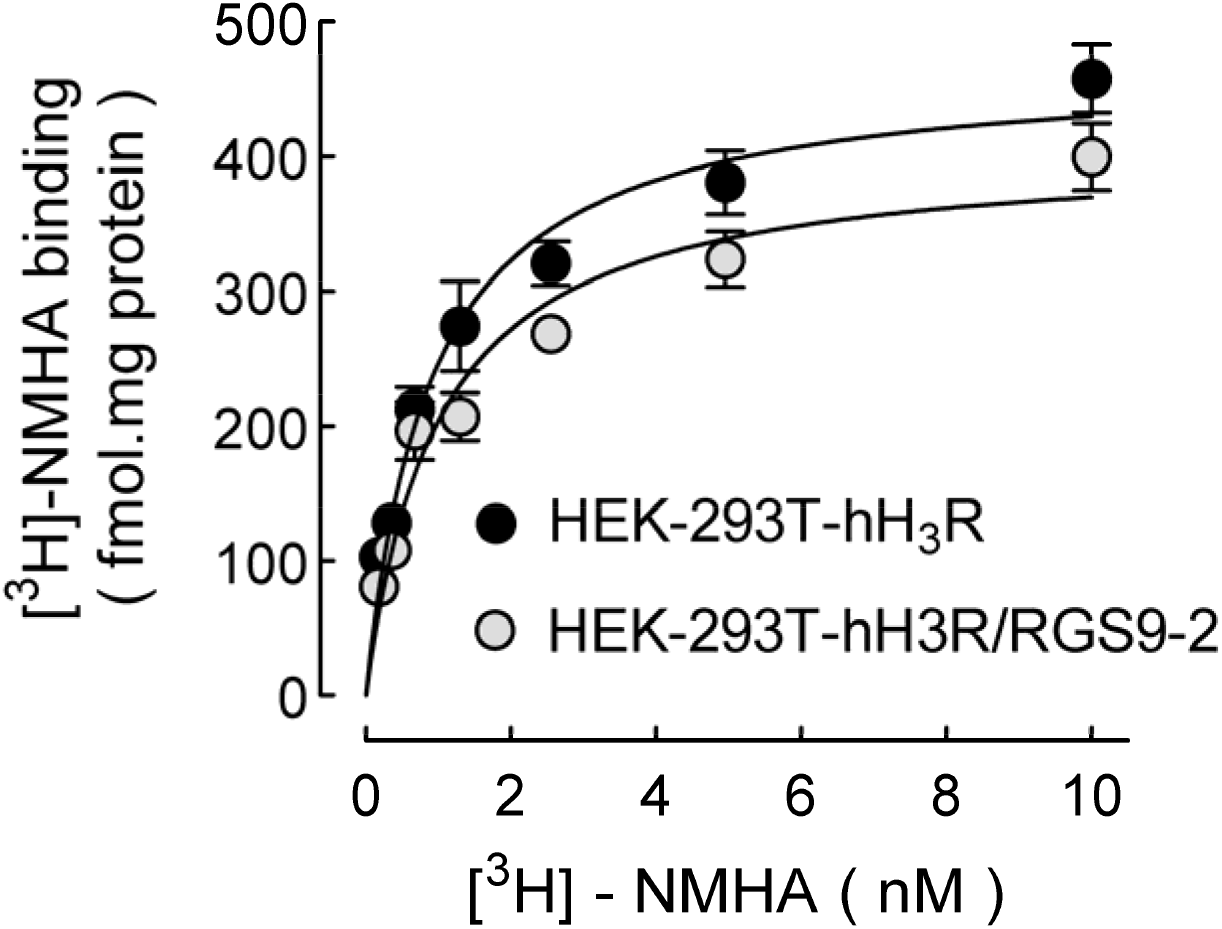
Saturating [^3^H]-NMHA binding to membranes from HEK293T-hH_3_R or HEK293T-hH_3_R/hRGS9-2 cells. Membranes were incubated with the indicated concentrations of [^3^H]-NMHA, and specific receptor binding was determined by subtracting the binding in the presence of 10 μM histamine from total binding. Points are means ± SEM from triplicate determinations in a single experiment, repeated a further 3 or 4 times. The line drawn is the best fit to a hyperbola. Best-fit values for the equilibrium dissociation constant (Kd) and maximal binding (Bmax) are given in the text.

### 3.2 Effect of hRGS9-2 on hH_3_R-stimulated [^35^S]-GTPγS binding

The capability of GPCRs to activate G proteins can be evaluated by [^35^S]-GTPγS binding to cell membranes. hH_3_R activation with the agonist immepip stimulated [^35^S]-GTPγS binding in membranes from either HEK293T-hH_3_R or HEK293T-hH_3_R/hRGS9-2 cells (Figure 3A). Neither the maximal response (E_max_) nor the agonist potency (pEC_50_) was different between the two cell types (Table 1). Furthermore, basal [^35^S]-GTPγS binding was not statistically different (377 ± 81 and 355 ± 93 fmol/mg protein for HEK293T-hH_3_R or HEK293T-hH_3_R/hRGS9-2 cells, respectively, 4 experiments, *p* = 0.865).

**Figure 3.**
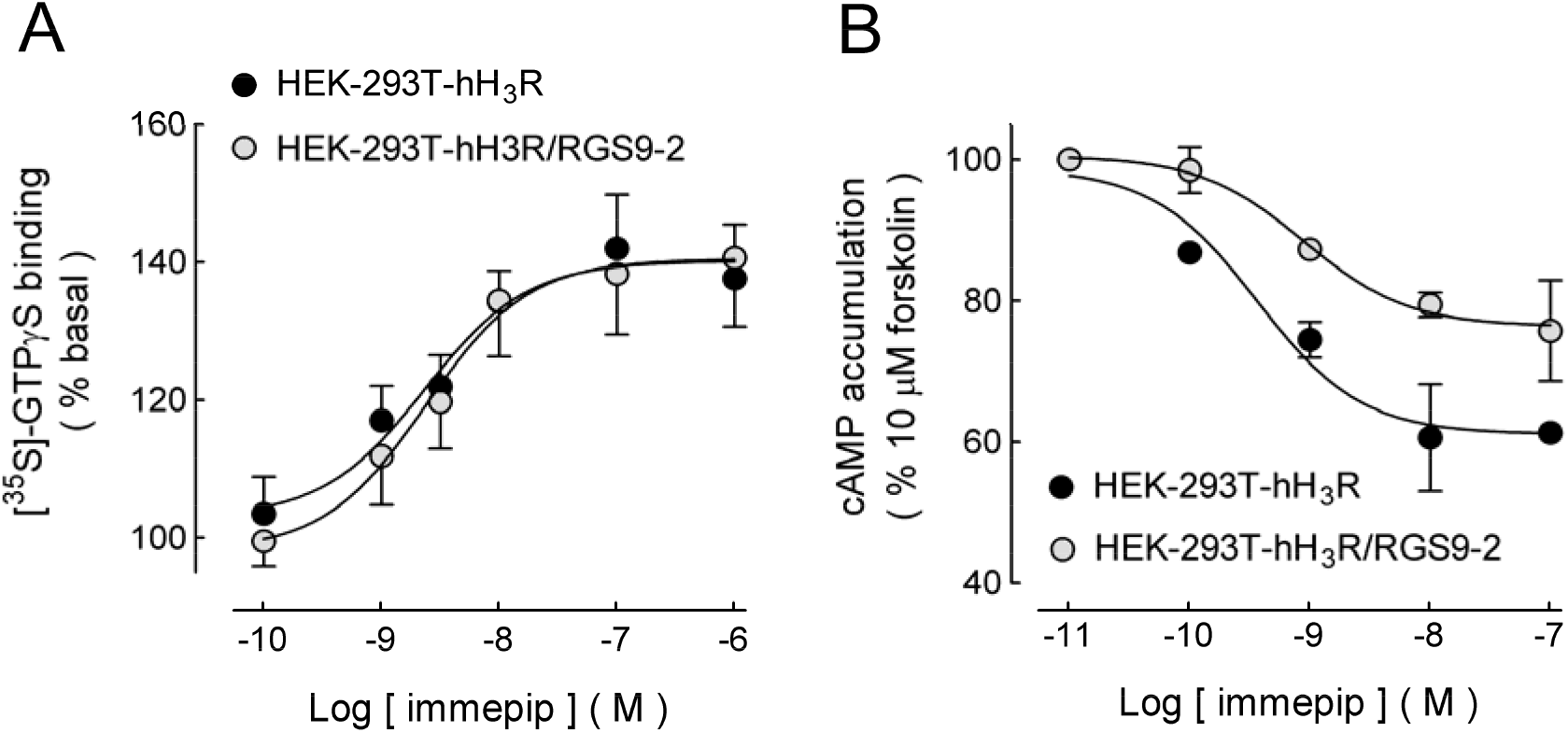
Stimulation of [^35^S]-GTPγS binding and inhibition of forskolin-induced cAMP accumulation elicited by H_3_R activation. A. [^35^S]-GTPγS binding. Membranes from HEK293T-hH_3_R or HEK293T-hH_3_R/hRGS9-2 cells were incubated (60 min) with 100 pM [^35^S]-GTPγS in the presence and absence of the indicated concentrations of the H_3_R agonist immepip. Values are expressed as percentage of basal [^35^S]-GTPγS binding after subtraction of nonspecific binding and correspond to the means ± SEM from three replicates from a representative experiment repeated a further 3 times with similar results. The line drawn is the best fit to a logistic equation. Best-fit values for maximal stimulation (E_max_) and pEC_50_ (−log_10_ EC_50_) are given in Table 1. B. Inhibition of forskolin-induced cAMP accumulation. HEK293T-hH_3_R or HEK293T-hH_3_R/hRGS9-2 cells were pre-incubated (15 min) in Krebs-Ringer-Henseleit solution containing 1 mM 3-isobutyl-1-methylxantine (IBMX) and then exposed for 30 min to forskolin (10 μM). When required immepip was added 5 min before forskolin. Values are expressed as percentage of the response to forskolin, after basal subtraction, and correspond to the means ± SEM from four replicates from a representative experiment, repeated a further 10 times. The line drawn is the best fit to a logistic equation. Values for maximum inhibition and pIC_50_ (−log_10_ IC_50_) are compared in Table 1.

**Table 1.** Comparison of the stimulation of [^35^S]-GTPγS binding and inhibition of forskolin-induced cAMP accumulation elicited by hH_3_R activation in HEK293T-hH_3_R or HEK293T-hH_3_R/hRGS9-2 cells.

|  | HEK293T-hH<sub>3</sub>R | HEK293T-hH<sub>3</sub>R/hRGS9-2 | *p* |
| --- | --- | --- | --- |
| [$^{35}$S]-GTPγS binding | | | |
| *E*<sub>max</sub> (%) | 146.3 ± 4.4 | 150.0 ± 5.3 | 0.617 |
| pEC<sub>50</sub> | 8.57 ± 0.26 | 9.00 ± 0.33 | 0.345 |
| cAMP accumulation |  |  |  |
| *I*<sub>max</sub> (%) | 37.7 ± 5.1 | 19.2 ± 5.3 | 0.022 |
| pIC<sub>50</sub> | 9.60 ± 0.14 | 9.07 ± 0.29 | 0.126 |

Values are means ± s.e.m. from 4 experiments for [ $^{35}$ S]-GTPγS binding and 11 experiments for cAMP accumulation. The statistical analysis was performed with Student's *t* test.

### 3.3 Effect of hRGS9-2 on H_3_R-mediated inhibition of forskolin-stimulated cAMP accumulation

The H_3_R agonist immepip inhibited forskolin-stimulated cAMP accumulation in both HEK293T-hH_3_R and HEK293T-hH_3_R/hRGS9-2 cells (Figure 3B), but the maximal inhibition was significantly larger for HEK293T-hH_3_R cells (Table 1). In spite of a trend for reduced agonist potency in cells co-transfected with hRGS9-2, the values did not reach statistical significance (Table 1).

Basal cAMP accumulation was similar for both isoforms (1.24 ± 0.19 and 1.83 ± 0.32 pmol/well for HEKT293-hH_3_R and HEK293T-hH_3_R/RGS9-2 cells, respectively, *p* = 0.134; 11 experiments, Student’s *t* test). Furthermore, there was no significant difference in the stimulatory effect of forskolin (12.95 ± 2.05 and 15.70 ± 2.44 pmol/well, *p* = 0.398).

## 4. Discussion

H_3_Rs couple to the Gα_i/o_ family proteins and are abundantly co-expressed with the protein RGS9-2 by striatal medium spiny neurons. By using transiently transfected HEK-293T cells, we showed here that the over-expression of RGS9-2 had no effect on the hH_3_R expression levels and nor did in the H_3_R-mediated G protein activation, nonetheless reduced the signaling of Gα_i/o_ proteins.

RT-PCR assays indicated that HEK-293T-hH_3_R and HEK-293T-hH_3_R/hRGS9-2 cells transcribed the HRH3 gene to a similar extent. Although the performed PCR was not quantitative, the amplicons were similar for cycles 22 to 28. Western blot analysis corroborated the translation of the mRNA encoding the hH_3_R and the hRGS9-2 protein in both cells. Levels of hRGS9-2 mRNA and protein increased, although not significantly, in cells transfected only with the hH_3_R (Figure 1). This effect could be underlain by a modulatory feedback [9], because in the striatum of patients with Parkinson’s disease the increase in dopamine D_2_ receptors [12] is accompanied by RGS9-2 over-expression [13]. Nevertheless, in cells co-transfected with human H_3_R and RGS9-2, the protein levels of hRGS9-2 were more than 2 fold of those in cells transfected only with the hH_3_R, and [^3^H]-NAMH binding assays indicated that hRGS9-2 transfection did not affect hH_3_R levels or the affinity of the receptors for the labeled agonist, supporting an appropriate model for hRGS9-2 over-expression.

The first step in hH_3_R-mediated signaling, the activation of Gα_i/o_ proteins, was evaluated by [^35^S]-GTPγS binding assays in which neither the efficacy nor the potency of the agonist immepip was modified by the co-transfection of hRGS9-2. G protein activation results from allosteric conformational changes in the Gα subunit induced by agonist-occupied GPCRs [14]. RGS proteins enhance the GTPase activity of activated Gα subunits and it is therefore assumed that neither the receptor/G protein interaction nor G protein activation is affected by RGS proteins [15]. However, in neuroblastoma N2A cells RGS2 siRNA knockdown enhanced [^35^S]-GTPγS binding induced by the activation of dopamine D_2_ receptors [16]. This discrepancy may be related to different levels of membranal D_2_ receptors in N2A cells because RGS2 knockdown prevented receptor internalization induced by exposure to an agonist for 30 min, the period used in cAMP assays [16], whereas the hH_3_R requires a longer time to experience homologous desensitization [17].

In regard to the signaling mediated by Gα_i/o_ proteins, in N2A cells RGS2 siRNA knockdown augmented D_2_ receptor-mediated inhibition of cAMP accumulation [16], and in this work RGS9-2 over-expression in HEK-293T cells resulted in reduced efficacy of the hH_3_R to inhibit forskolin-induced cAMP accumulation, supporting that RGS proteins down-regulate the signaling of Gα_i/o_ proteins. Reduced signaling was also observed for the D_3_ receptor in HEK-293 cells upon RGS9-2 over-expression [18], although the extent of the effect was larger for the hH_3_R (49 % versus ~20 %). Interestingly, the effect of RGS9-2 appears receptor-specific because its over-expression in HEK-293 cells had no effect on the signaling of the D_2_ receptor, which belongs to the same family (D_2_-like) as the D_3_ receptor, whereas the signaling of both D_2_ and D_3_ receptors was reduced by RGS4 [18]. The distinctive regulation by RGS9-2 of the signaling of GPCRs coupled to Gα_i/o_ proteins could be explained by the differential participation of the receptors in the formation of protein complexes, in particular with Gβ5 and R7BP, that simultaneously interact with RGS9-2 and control its localization in the plasma membrane [9].

One clear limitation of this study is the experimental model, which cannot mimic the complexities of the natural conditions in neurons that endogenously express H_3_Rs, RGS9-2 and their interacting proteins. However, the possible modulation by RGS proteins of H_3_R function is virtually unexplored and this work may thus provide the basis for future studies.

## 5. Conclusion

The data presented herein indicate that the hRGS9-2 protein regulates the cell signaling elicited by the hH_3_R in a heterologous expression system, suggesting a similar scenario for striatal spiny GABAergic neurons that express high levels of H_3_Rs and RGS9-2.

## Abbreviations

CNS: Central Nervous System
GPCR: G protein-coupled receptor
H_3_R: histamine H_3_ receptor
hH_3_R: human histamine H_3_ receptor
RGS: regulator of GPCR signaling
hRGS9-2: human regulator of GPCR signaling 9-2

## Acknowledgements

We are grateful to Raúl González-Pantoja for his very excellent technical assistance. G. Nieto-Alamilla held a scholarship from the Mexico State Council for Science and Technology (Comecyt).

## Author contribution

G. N.-A. and J.-A. A.-M. designed the study; G. N.-A. and J. E.-S. performed the experiments; G. N.-A. and J.-A. A.-M. performed data analysis. G.N.-A. and J.-A. A.-M. wrote the manuscript. All three authors revised and approved the manuscript.

